# Development of a BiAD sensor for locus-specific detection of cellular histone acetylation dynamics by fluorescence microscopy

**DOI:** 10.1101/2025.02.10.637448

**Authors:** Anja R. Köhler, Nicole Gutekunst, Annika Harsch, Pavel Bashtrykov, Albert Jeltsch

## Abstract

Dynamic changes in histone acetylation play crucial roles during cellular differentiation and disease development. Here, we developed a Bimolecular Anchor Detector (BiAD) sensor for the detection of locus-specific changes in histone acetylation in living cells by fluorescence microscopy. For this, we used the BRD9 bromodomain cloned as tandem double-domain (2xBRD9-BD) as reader of histone acetylation. It was integrated it into a dual-color BiAD chassis that was previously described [Köhler et al., 2024, Cell Rep Methods 4(4):100739]. We identified the gene body of TTC34 as potential target for our sensor, because it contains dense histone acetylation and 392 local sequence repeats. Using a binding deficient mutant of 2xBRD9-BD as negative control, we established successful readout of histone acetylation at the TTC34 locus. A single domain reader did not function indicating the requirement for the double reader to enhance affinity and specificity of the chromatin interaction via avidity effects. With this sensor, we could detect dynamic changes of histone acetylation at the TTC34 locus after treatment of cells with the histone deacetylase inhibitor Trichostatin A for 6 hours indicating the applicability of this sensor for single-cell epigenome studies.

## Introduction

Histone lysine acetylation is an integral part of the epigenome and central to the epigenetic control of gene transcription [1, 2]. Chromatin modifications are bound by special reader proteins which trigger downstream signaling events [3]. The bromodomain (BD) has been found in many chromatin-associated proteins and enzymes, and shown to function as a general reader domain for acetyllysine residues on histones and other proteins [4, 5]. Based on this property, BD containing proteins have been fused to fluorophores and used for global detection of histone acetylation in cells since many years and the corresponding sensor systems have been continuously improved [6-9]. However, all the so far established systems cellular sensor systems do not permit detection of histone acetylation at defined genomic loci. Hence global histone acetylation changes observed with these sensors in living cells cannot be connected with individual genes which limits the applicability of these sensors. On the contrary, analysis of histone acetylation by ChIP-seq (and variants thereof) requires the lysis of cells prevailing the direct association of specific acetylation patterns with cellular phenotypes. Hence, novel techniques for detection of histone acetylation with locus resolution in living cells are needed.

During the last years, we developed bimolecular anchor detector (BiAD) sensors to overcome this limitation and allow for locus-specific readout of epigenome modifications in living cells [10, 11]. In principle, the BiAD sensors comprise a programmable DNA-binding anchor module for locus-specific targeting of the sensor, which is a single-guide RNA (sgRNA)/dCas9 (deactivated Cas9) complex in the most recent applications. This module is combined with chromatin reader domains used as detector modules for the recognition of defined chromatin modifications at the genomic target locus. Both modules are fused to complementary parts of a split fluorophore, which leads to bimolecular fluorescence complementation when both modules bind in close spatial proximity indicating that the target locus contains the chromatin modification of interest [10]. In a recent work, the 1^st^ generation BiAD sensors (described in [10]) were massively improved [11] in three aspects: 1) To enhance the sensitivity of BiAD sensors, a signal amplification using the 10xSunTag was implemented which enabled the recruitment of multiple detector modules to a single sgRNA/dCas9 binding site. 2) The epigenetic readout is combined with a second YPet fluorophore which is recruited by the sgRNA and labels the locus of interest. The resulting dual-color BiAD sensors allow to detect the BiAD signal specifically at the locus of interest, which led to a strong improvement of signal-to-noise ratio, because background fluorescence fluctuation and non-specific protein aggregation could be excluded from the analysis. 3) As a third improvement, it was demonstrated that double reader domains generally perform better in BiAD applications, finally leading to a system that allowed the detection of DNA methylation, as well as di- and trimethylation of H3K9, H3K27 and H3K36 at endogenous loci with a minimum of 45 local repeats.

BiAD sensors had been used to detect the dynamics of DNA methylation and H3K9 methylation upon DNA or protein methyltransferase KO or KD, or after treatment of cells with a methyltransferase inhibitor [10, 11]. Moreover, elevated H3K9 methylation signals could be detected at the inactive X-chromosome [11], locus-specific reductions of H3K9me3 at mouse major satellite repeats were found in p53 deficient cells [12], and changes in human α-satellite repeat DNA methylation were detected depending on the density of culturing [13]. All these biological applications demonstrate the wide applicability of the BiAD sensors, even at their current state of development being able to detect epigenome modifications only at regions with a high to moderate level of local sequence repetitions. However, so far no BiAD sensor was available for active chromatin modifications, like H3K4me3 or histone acetylation. It was the aim of this work to fill this gap by developing and validating a functional dual-color BiAD sensor for the detection of histone acetylation at defined genomic loci in living cells. For this, an acetyllysine reader domain needed to be developed, integrated into the existing dual-color BiAD chassis and the functionality of the BiAD sensor for histone acetylation needed to be validated. Finally, we aimed for a demonstration of the application of the novel BiAD sensor to detect dynamic changes of histone acetylation at a defined target locus in living cells.

## Materials and Methods

### Cloning of the acetyllysine detector domains

The bromodomain of BRD9 (Uniprot Q9H8M2-1, amino acids 14-134) was amplified from HEK293 cDNA and cloned into the BiAD detector-IFP2.0C vector [11] using Gibson Assembly. To create a binding deficient detector mutant, the conserved tyrosine residue Y57 was substituted with alanine via site directed mutagenesis [14], because the Y-to-A substitution of the corresponding residue in BRD2 (Y113, Uniprot P25440-1) was shown to disrupt H4K12ac binding [6, 8, 15]. The double domain construct, in which both domains are separated by a 12 amino acid linker containing an SV40 large T-antigen monopartite NLS was cloned as described [11].

### Cell culture

Human embryonic kidney cells (HEK293, RRID: CVCL_0045) were sourced from DSMZ (Braunschweig, Germany). The cells were cultured in DMEM (Dulbecco’s Modified Eagle Medium) supplemented with 10% fetal bovine serum (FBS), 1% penicillin/streptomycin, and 4 mM L-glutamine, and maintained at 37 °C with 5% CO_2_ in a humidified incubator (BINDER). To sustain a confluency of 70-90%, the cells were sub-cultured at a 1:7 to 1:10 ratio every 2 to 3 days. For this, the cells were washed with PBS (lacking CaCl_2_ and MgCl_2_), followed by addition of trypsin-EDTA solution (Sigma), and incubation at 37 °C until the cells detached. Afterwards, the cells were resuspended in fresh culture medium and split at the required ratio. For cryopreservation, cells were pelleted at 300 g for 5 minutes, resuspended in freezing medium (90% FBS and 10% DMSO), and gradually frozen to -80 °C.

### Transfections

For transient transfections, HEK293 cells were seeded on glass cover slips in a 6-well plate at a density of 200,000 cells per well one day before transfection. Right before transfection, the medium was exchanged with 1.5 mL fresh culture medium supplemented with 30 nM biliverdin which serves as the chromophore for IFP2.0. Cells were transfected with 400 ng sgRNA, 100 ng dCas9-10xSunTag, 300 ng scFv-IFP2.0N, 300 ng BRD9BD-IFP2.0C/2xBRD9BD-IFP2.0C and 400 ng MCP-YPet using polyethylenimine (PEI) at a ratio of 1:3 as the transfection reagent. Cells were fixed with 4% paraformaldehyde 24 h after transfection as described in [11].

### HDAC inhibitor treatment

HEK239 cells were transfected as described in the previous section and treated with 5 µM Trichostatin A (TSA, PubChem compound 444732) (Sigma) or an equivalent amount of DMSO for six or three hours before fixation.

### Fluorescence microscopy and image analysis

The fixed cell samples were imaged using a confocal laser scanning microscope (LSM710 or LSM980 Airyscan 2, Carl Zeiss) using the image acquisition settings described [11]. The settings were kept constant within the same experiment. The stacks were superimposed to generate maximum intensity projections using the ZEN 3.0 SR (black edition) software. Image export was performed using the ZEN 3.0 (blue edition) software. The exemplary images in each figure were exported with identical brightness and contrast settings.

Images were analyzed using a custom FIJI (ImageJ 1.54b) [16] macro without adjustments of contrast and brightness as described [11]. In brief, two intensity thresholds were set manually for each cell to define the nucleus and spots as regions of interest (ROIs) based on the signal in the YPet channel. The mean intensity of the ROIs was measured across all channels. For quantitative analysis, the average nuclear background signal intensity was subtracted from the spot intensities in each channel. The BiAD spot intensity (IFP2.0 channel) was then normalized to the corresponding YPet spot intensity and the normalized BiAD signals from all spots within a single cell were averaged and are represented as a dot in the boxplots. Boxplots were created using the Seaborn Python data visualization library [17, 18].

## Results

BiAD sensors fill an important gap in the available technologies by allowing the detection of epigenome modifications with locus specificity in single living cells by fluorescence microscopy [10, 11]. The most recently developed dual-color BiAD sensors [11] combine two fluorescence signals and two orthogonal signal amplification systems (Figure 1A). The target locus is bound by a sgRNA/dCas9 complex and its location is indicated by the recruitment of several YPet fluorophores to the sgRNA via the MS2 system [19]. The readout of the epigenome modification is based on reconstitution of the N- and C-terminal parts of the split IFP2.0 fluorophore [20, 21]. The N-terminal part of IFP2.0 is fused to a single-chain variable fragment (scFv) antibody domain binding to the GCN4 peptide epitope which is present in 10 copies in the so called 10xSunTag fused to the dCas9 [22]. The C-terminal part of IFP2.0 is fused to an appropriate epigenome reader domain for detection of the epigenome modification of interest at the target locus. In the past years, BiAD sensors have been developed for the locus-specific detection of DNA methylation as well as H3K9, K27 and K36 methylation, but so far, a detector for an active chromatin mark is missing in the BiAD toolbox. Here we aimed to develop and validate such a novel tool for the detection of histone acetylation.

**Figure 1:**
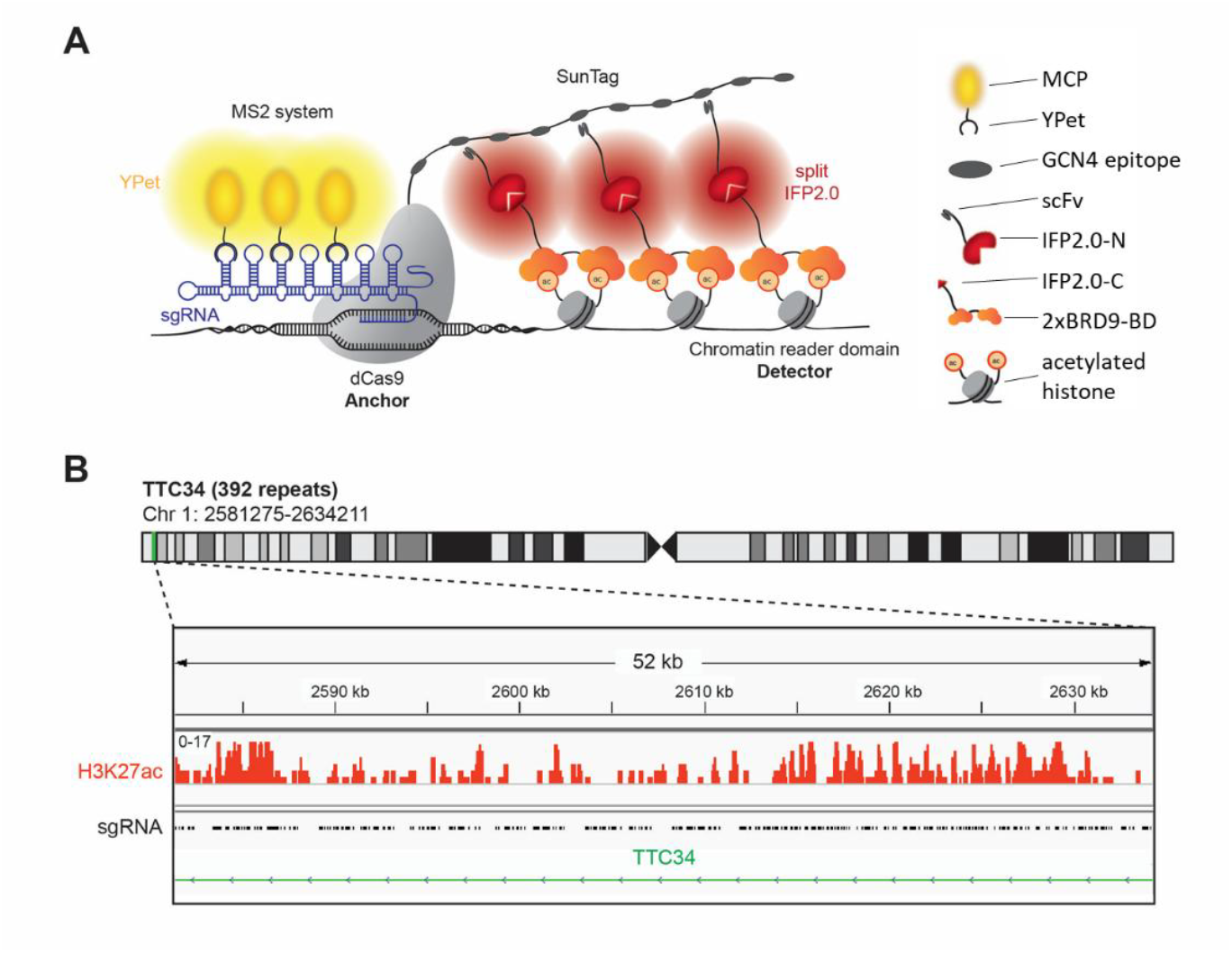
Development of a dual-color BiAD sensor for detection of histone acetylation. **A)** Schematic representation of the sensor design. The target locus is visualized by a full-length fluorophore (YPet), recruited by an MS2 scaffold of the sgRNA in complex with the anchor module (dCas9). One part of the split fluorophore (IFP2.0) is recruited to the SunTag amplification system fused to dCas9 by a single-chain variable fragment (scFv) antibody domain. To detect acetylated histones, a double BRD9 bromodomain (2xBRD9-BD) is fused to the second part of the split IFP2.0 fluorophore. If histone acetylation (Kac) is present at the target locus, the detector module can bind and bring the second part of the split IFP2.0 in close spatial proximity to the first one leading to reconstitution of the split IFP2.0 and appearance of fluorescent BiAD signal. **B)** Identification of a repetitive genomic region containing histone acetylation, which can be used for the validation of the novel detector module. The gene body of the TTC34 gene contains 392 repeats of the sgRNA binding sites (indicated in the black trace) and it is modified with H3K27ac (indicated in the orange trace).

### Identification of a target region for histone acetylation detection with the dual-color BiAD sensor

Despite their enhanced sensitivity, even the most developed 2^nd^ generation dual-color BiAD sensors require the presence of at least 45 local DNA repeats at the target region allowing for the binding of several sgRNA/dCas9 complexes to generate visible BiAD signals [11]. This requirement entailed a critical problem, because repetitive sequences in the human genome usually are epigenetically silenced. Therefore, there were no obvious candidate regions containing histone acetylation at local repeats that could be used as a target for the development of a BiAD sensor for histone acetylation. To search for a suitable target region, we resorted to a list of repetitive regions containing potential sgRNA binding sites in the human genome [19] which was compared with available ChIP-seq data for HEK293 cells [23]. HEK293 cells were selected for the development of the BiAD sensors, because of their easy handling and high transfectability. This comparison identified the gene body of TTC34 as potential target for the measurement of histone acetylation with a BiAD sensor, as it contains 392 local repeats of a CCACAGGTGAGCATC sequence [19] and dense histone acetylation (Figure 1B).

### Integration of a functional acetyllysine reader domain into the dual-color BiAD chassis

As described above, bromodomains have been identified as readers of acetyllysine [4, 5]. More specifically, the bromodomain of BRD9 was identified in peptide spot binding experiments as pan-histone acetylation detector [24]. Based on different previous studies from other labs [9, 25-27] and our own data [11, 28], we expected that the binding affinity and specificity of the BRD9-BD as detector module toward histone acetylation would be improved by using two BDs fused in a tandem arrangement into a double domain, called 2xBRD9-BD. The BRD9-BD was cloned from human cDNA and the single and double domain integrated into the existing dual-color BiAD sensor addressing the TTC34 repeats [11]. To generate a binding deficient BRD9-BD as negative control, we mutated the conserved tyrosine residue Y57 in the BRD9-BD and 2xBRD9-BD context to alanine, because the corresponding Y113A mutation in bromodomain 1 (BD1) of the BRD2 protein (Supplementary Figure 1) was shown to disrupt H4K12ac binding [6, 8, 15].

HEK293 cells were transfected with all components of the dual-color BiAD sensor for histone acetylation detection at the TTC34 target locus with either the wildtype 2xBRD9BD detector module (WT) or the Y57A mutant. The cells were inspected by fluorescence microscopy 24 h after transfection. They showed strong colocalization of the BiAD signal (IFP2.0) with the locus marker fluorophore (YPet) for the WT detector, but not for the binding-deficient Y57A mutant (Figure 2A). A quantitative analysis revealed a highly significant increase in BiAD signal at the target loci in all three independent experiments with an intact (WT) detector modules, but not with the mutant detector modules (Figure 2B and 2C). These data validate the function of the novel BiAD sensor for detection of histone acetylation. Similar experiments with the BRD9 single BD did not results in visible or statistically significant BiAD signal increase at target loci with the WT reader domain when compared with the inactive mutant, indicating that the single BD BiAD reader is not functional (Supplementary Figure 2). Hence, double reader domains improve the functionality of the BiAD sensor though avidity effects as previously observed for the HP1ß-chromodomain, DNMT3A-PWWP domain and CBX7-chromodomain readers of H3K9, H3K36 and H3K27 di- and trimethylation [11].

**Figure 2:**
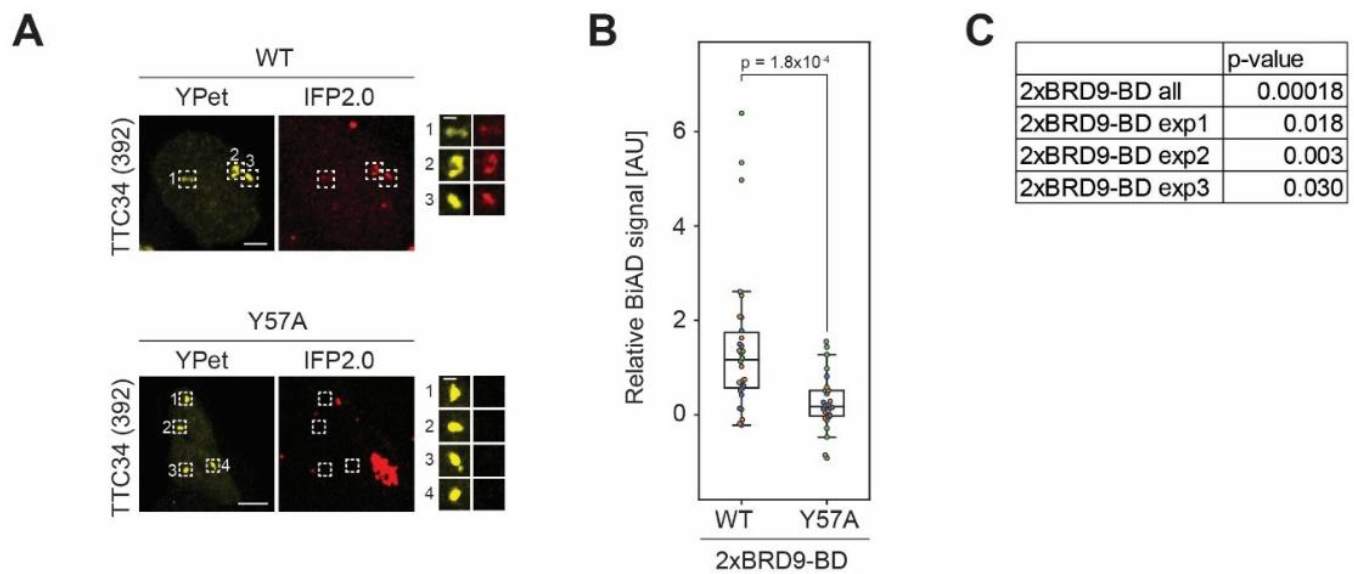
Validation of the BRD9-BD double domain (2xBRD9-BD) for histone acetylation readout at the TTC34 locus. HEK293 cells were transfected with all components of the dual-color BiAD sensor for histone acetylation detection at the TTC34 target locus with either the wildtype 2xBRD9-BD detector (WT) or a binding-deficient mutant (Y57A). **A)** Exemplary fluorescence microscopy images showing strong BiAD signal (IFP2.0) at the marker fluorophore (YPet) spots with the for the WT detector, but not for the binding-deficient Y57A mutant. Scale bars are 5 μm and 1 μm for the magnified images. **B)** Boxplot showing the relative BiAD signals of three independent experiments (depicted in orange, blue and green). Significance was determined via a two-tailed, unpaired t-test. **C)** P-values between WT and Y57A for all three combined data sets as well as each for each of the individual experiments. See also Supplementary Figure 2.

### Application of the novel BiAD reader to detect dynamic changes of histone acetylation in living cells

Next, we were aiming to apply the newly developed BiAD sensor for histone acetylation at the TTC34 locus to detect dynamic changes of this modification in living cells. To this end, HEK293 cells were transfected with the plasmids encoding the sensor components and either treated with TSA for 3 and 6 hours or treated with DMSO as negative control. TSA is a well-established inhibitor of class I and II histone deacetylases [29]. When applied to the cell culture medium it leads to a strong increase in histone acetylation. Comparison of the BiAD signals of the control sample and cells treated with TSA for 6 hours indicated clear and statistically highly significant increase of the BiAD signal in the case of the intact sensor system, but not with the system including the binding deficient Y57A BD (Figure 3 and Supplemental Figure 3). Moreover, this experiment confirmed the detection of histone acetylation at the target loci with the WT but not with the mutant sensor (Figure 3 and Supplemental Figure 3B).

**Figure 3:**
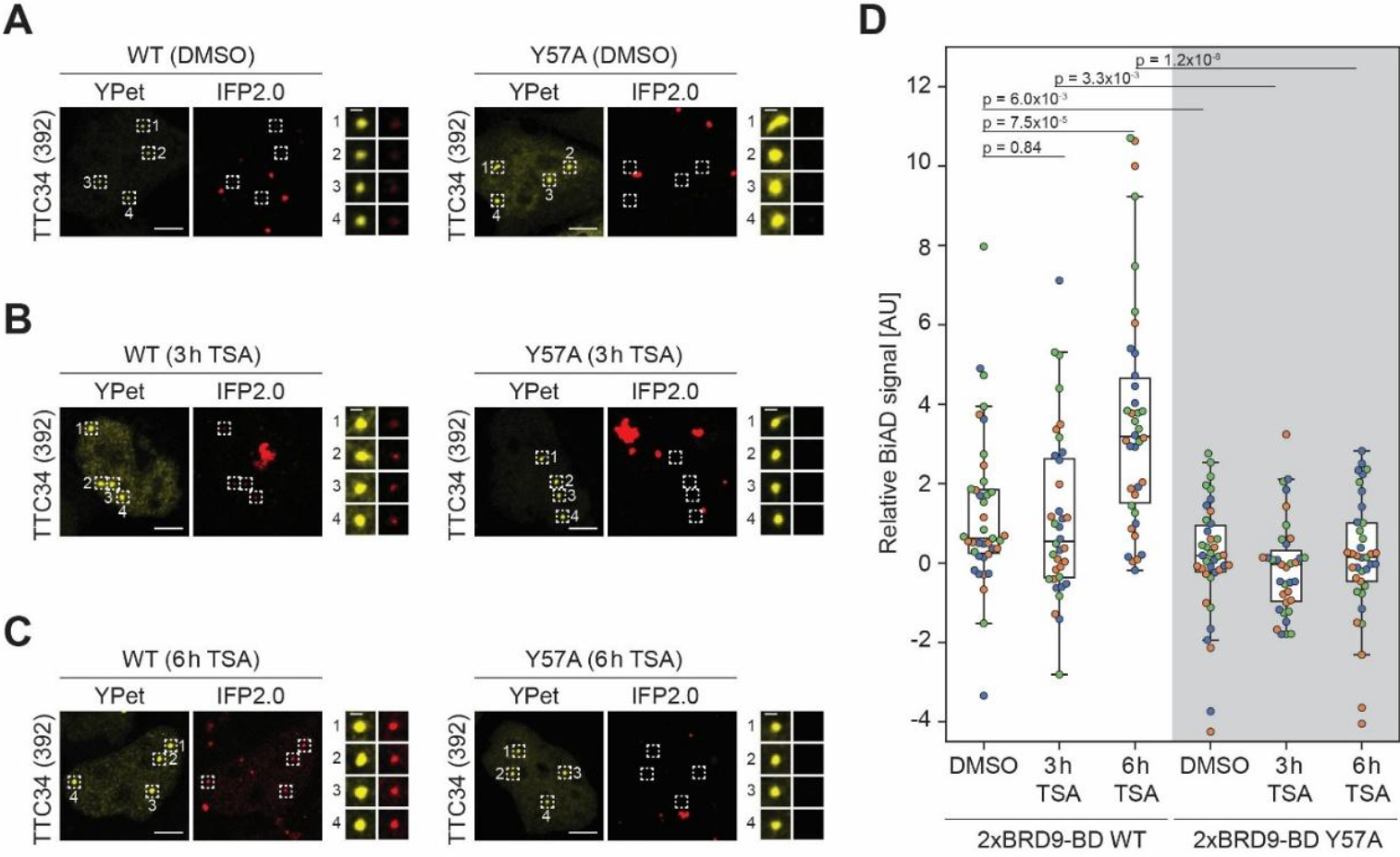
Visualization of locus-specific gain in histone acetylation upon TSA treatment. HEK293 cells were transfected with all components of the dual-color BiAD sensor for histone acetylation detection at the TTC34 target locus with either the wildtype 2xBRD9-BD detector (WT) or a binding-deficient mutant (Y57A). Cells were either mock treated (DMSO) or treated with TSA for 3 h or 6 h before fixation. **A)-C)** Exemplary fluorescence microscopy images showing colocalization of the BiAD signal (IFP2.0) with the marker fluorophore (YPet) for the WT detector, but not for the binding-deficient Y57A mutant. The cells were either treated with DMSO (A), for for 3 (B) or 6 h with TSA (C). **D)** Boxplot showing the relative BiAD signals of three independent experiments (depicted in orange, blue and green). The significance of differences of the BiAD signals was determined via a two-tailed, unpaired t-test. P-values are indicated in the boxplot. The WT BiAD sensor showed an increase in the IFP2.0 signal in cells after 6 h treatment with TSA compared to the DMSO negative control. Additional pairwise p-values are provided in Supplementary Figure 3B.

## Discussion

In this work we describe a dual-color BiAD sensor for the detection of histone acetylation. Our data demonstrate that active chromatin modification can be detected by BiAD sensors which complements the toolkit of available BiAD sensors. The results of our study confirm the modularity of BiAD sensors [10, 11] suggesting that additional readers for even more epigenome modifications could be added in a similar fashion.

However, while demonstrating the clear capability of locus-specific detection of histone acetylation at the TTC34 gene body, the BiAD sensor developed here suffers from a similar problem as its predecessors described in Köhler et al. (2024) [11]. This is illustrated by the observation in Figure 2 that even in the system using the WT 2xBRD9-BD detector module many cells apparently were BiAD negative. While one cannot rule out cellular heterogeneity as a reason for this variability, there is an important technical issue to be considered, which is that the BiAD sensor used here depends on the co-transfection of 5 individual plasmids. These plasmids encode dCas9-10xSunTag (plasmid 1), MCP-YPet (plasmid 2), sgRNAs (plasmid 3), scFv-IFP2 fused to the N-terminal part of IFP2.0 (plasmid 4), and the fusion of reader domain with the C-terminal part of IFP2.0 (plasmid 5). Among them, successful transfection of plasmids 1-3 can be validated by the appearance of the YPet signal at the target locus, allowing to restrict the analysis on cells containing these 3 plasmids. However, it remained invisible if target cells really contained the plasmids 4 and 5, and, given the limitations of co-transfections, it is likely that some of the apparently BiAD negative cells observed with the WT 2xBRD9-BD BiAD sensor in reality were “false negatives”, because they did not contain one of these plasmids, and hence did not harbor a functional BiAD sensor. This problem could be overcome in future work by the generation of stable cell lines expressing the BiAD components encoded by plasmids 4 and 5 or by strategically combining multiple BiAD components on the same plasmid to make successful transfection of all plasmids visible.

Such a conceptional improvement should considerably increase the dynamic range between the signals observed with WT and binding-deficient mutant detector BiAD sensors. This would directly enhance the sensitivity of the sensor. Whether such improvement of the sensitivity will be sufficient to reach single locus resolution remains to be tested experimentally. Moreover, the elimination of false negative cells from the analysis would facilitate true single cell analysis of the BiAD signals by removing the need to analyze entire cell populations for statistical purpose in order to draw conclusions about individual cells. In such experimental setting, single living cells could be tracked over time to directly correlate changes in the locus-specific epigenomic landscape to alterations in the cellular phenotype.

## Acknowledgements

The authors gratefully acknowledge the Technology Platform “Cellular Analytics” of the Stuttgart Research Center Systems Biology for their support and assistance in this work. The initial parts of this work have been supported by the Baden-Württemberg Stiftung gGmbH (EpiSensor, ID07 to AJ). The funder had no role in the design of the study and collection, analysis, and interpretation of data and in writing the manuscript which should be declared.

## Author contributions

ARK and AJ devised the study. ARK conducted all experiments with the help of NG and AH. ARK, PB, and AJ were involved in data analysis. PB and AJ supervised the work. ARK and AJ wrote the draft article and prepared the figures. AJ acquired funding. All authors were involved in preparing the final manuscript.

### Data and code availability

All analyzed data are included in the published article and its supplementary information. The plasmids encoding the 2xBRD9-BD fused to the C-terminal part of IFP2.0 were deposited with Addgene. The ImageJ macro used for image analysis is available at Figshare (https://doi.org/10.6084/m9.figshare.23592894).

## References

1. Allis, C. D. & Jenuwein, T. (2016) The molecular hallmarks of epigenetic control, Nat Rev Genet. 17, 487–500.

2. Shvedunova, M. & Akhtar, A. (2022) Modulation of cellular processes by histone and non-histone protein acetylation, Nat Rev Mol Cell Biol. 23, 329–349.

3. Patel, D. J. & Wang, Z. (2013) Readout of epigenetic modifications, Annu Rev Biochem. 82, 81–118.

4. Marmorstein, R. & Zhou, M. M. (2014) Writers and readers of histone acetylation: structure, mechanism, and inhibition, Cold Spring Harb Perspect Biol. 6, a018762.

5. Jain, A. K. & Barton, M. C. (2017) Bromodomain Histone Readers and Cancer, J Mol Biol. 429, 2003–2010.

6. Kanno, T., Kanno, Y., Siegel, R. M., Jang, M. K., Lenardo, M. J. & Ozato, K. (2004) Selective recognition of acetylated histones by bromodomain proteins visualized in living cells, Mol Cell. 13, 33–43.

7. Sasaki, K., Ito, T., Nishino, N., Khochbin, S. & Yoshida, M. (2009) Real-time imaging of histone H4 hyperacetylation in living cells, Proc Natl Acad Sci U S A. 106, 16257–62.

8. Ito, T., Umehara, T., Sasaki, K., Nakamura, Y., Nishino, N., Terada, T., Shirouzu, M., Padmanabhan, B., Yokoyama, S., Ito, A. & Yoshida, M. (2011) Real-time imaging of histone H4K12-specific acetylation determines the modes of action of histone deacetylase and bromodomain inhibitors, Chem Biol. 18, 495–507.

9. Sanchez, O. F., Mendonca, A., Carneiro, A. D. & Yuan, C. (2017) Engineering Recombinant Protein Sensors for Quantifying Histone Acetylation, ACS sensors. 2, 426–435.

10. Lungu, C., Pinter, S., Broche, J., Rathert, P. & Jeltsch, A. (2017) Modular fluorescence complementation sensors for live cell detection of epigenetic signals at endogenous genomic sites, Nat Commun. 8, 649.

11. Kohler, A. R., Hausser, J., Harsch, A., Bernhardt, S., Haussermann, L., Brenner, L. M., Lungu, C., Olayioye, M. A., Bashtrykov, P. & Jeltsch, A. (2024) Modular dual-color BiAD sensors for locus-specific readout of epigenome modifications in single cells, Cell Rep Methods. 4, 100739.

12. Panatta, E., Butera, A., Mammarella, E., Pitolli, C., Mauriello, A., Leist, M., Knight, R. A., Melino, G. & Amelio, I. (2022) Metabolic regulation by p53 prevents R-loop-associated genomic instability, Cell Rep. 41, 111568.

13. Brenner, L. M., Meyer, F., Yang, H., Kohler, A. R., Bashtrykov, P., Guo, M., Jeltsch, A., Lungu, C. & Olayioye, M. A. (2024) Repeat DNA methylation is modulated by adherens junction signaling, Commun Biol. 7, 286.

14. Jeltsch, A. & Lanio, T. (2002) Site-directed mutagenesis by polymerase chain reaction, Methods in molecular biology (Clifton, NJ). 182, 85–94.

15. Umehara, T., Nakamura, Y., Jang, M. K., Nakano, K., Tanaka, A., Ozato, K., Padmanabhan, B. & Yokoyama, S. (2010) Structural basis for acetylated histone H4 recognition by the human BRD2 bromodomain, J Biol Chem. 285, 7610–8.

16. Schindelin, J., Arganda-Carreras, I., Frise, E., Kaynig, V., Longair, M., Pietzsch, T., Preibisch, S., Rueden, C., Saalfeld, S., Schmid, B., Tinevez, J. Y., White, D. J., Hartenstein, V., Eliceiri, K., Tomancak, P. & Cardona, A. (2012) Fiji: an open-source platform for biological-image analysis, Nat Methods. 9, 676–82.

17. Waskom, M. (2021) seaborn: statistical data visualization, Journal of Open Source Software. 6, 3021.

18. Hunter, J. D. (2007) Matplotlib: A 2D Graphics Environment, Computing in Science & Engineering. 9, 90–95.

19. Ma, H., Tu, L.-C., Naseri, A., Chung, Y.-C., Grunwald, D., Zhang, S. & Pederson, T. (2018) CRISPR-Sirius: RNA scaffolds for signal amplification in genome imaging, Nature Methods. 15, 928–931.

20. Tchekanda, E., Sivanesan, D. & Michnick, S. W. (2014) An infrared reporter to detect spatiotemporal dynamics of protein-protein interactions, Nature Methods. 11, 641–644.

21. Chen, M., Yan, C., Ma, Y. & Zhang, X.-E. (2021) A tandem near-infrared fluorescence complementation system with enhanced fluorescence for imaging protein-protein interactions in vivo, Biomaterials. 268, 120544.

22. Tanenbaum, M. E., Gilbert, L. A., Qi, L. S., Weissman, J. S. & Vale, R. D. (2014) A protein-tagging system for signal amplification in gene expression and fluorescence imaging, Cell. 159, 635–46.

23. Broche, J., Kungulovski, G., Bashtrykov, P., Rathert, P. & Jeltsch, A. (2021) Genome-wide investigation of the dynamic changes of epigenome modifications after global DNA methylation editing, Nucleic Acids Res. 49, 158–176.

24. Filippakopoulos, P., Picaud, S., Mangos, M., Keates, T., Lambert, J. P., Barsyte-Lovejoy, D., Felletar, I., Volkmer, R., Muller, S., Pawson, T., Gingras, A. C., Arrowsmith, C. H. & Knapp, S. (2012) Histone recognition and large-scale structural analysis of the human bromodomain family, Cell. 149, 214–31.

25. Villasenor, R., Pfaendler, R., Ambrosi, C., Butz, S., Giuliani, S., Bryan, E., Sheahan, T. W., Gable, A. L., Schmolka, N., Manzo, M., Wirz, J., Feller, C., von Mering, C., Aebersold, R., Voigt, P. & Baubec, T. (2020) ChromID identifies the protein interactome at chromatin marks, Nat Biotechnol. 38, 728–736.

26. Veggiani, G., Villaseñor, R., Martyn, G. D., Tang, J. Q., Krone, M. W., Gu, J., Chen, C., Waters, M. L., Pearce, K. H., Baubec, T. & Sidhu, S. S. (2022) High-affinity chromodomains engineered for improved detection of histone methylation and enhanced CRISPR-based gene repression, Nature communications. 13, 6975.

27. Tekel, S. J., Vargas, D. A., Song, L., LaBaer, J., Caplan, M. R. & Haynes, K. A. (2018) Tandem Histone-Binding Domains Enhance the Activity of a Synthetic Chromatin Effector, ACS Synth Biol. 7, 842–852.

28. Mauser, R., Kungulovski, G., Keup, C., Reinhardt, R. & Jeltsch, A. (2017) Application of dual reading domains as novel reagents in chromatin biology reveals a new H3K9me3 and H3K36me2/3 bivalent chromatin state, Epigenetics & chromatin. 10, 45.

29. Dokmanovic, M., Clarke, C. & Marks, P. A. (2007) Histone deacetylase inhibitors: overview and perspectives, Mol Cancer Res. 5, 981–9.

