## Supplemental Information for "Development of a BiAD sensor for locus-specific detection of cellular histone acetylation dynamics by fluorescence microscopy"

### Supplementary Information

#### Supplementary Figures

```
BRD9-BD   : PHGFFAFPVTD------AIAPGYSMIIKHPMDFGTMKDKIVANEYKSVTEFKADFKLMCDNAMTYNRPDTVYY : 107
BRD2-BD1  : WKHQFAWPFRQPVDAVKLGLPDYHKIIKQPMDMGTIKRRLENNYYWAASECMQDFNTMFTNCYIYNKPTDDIV : 163
BRD2-BD2  : KHAAYAWPFYKPVDASALGLHDYHDIIKHPMDLSTVKRKMENRDYRDAQEFAADVRLMFSNCYKYNPPDHDVV : 236
```

**Supplementary Figure 1: Sequence alignment of the BD of BRD9 with the BD1 and BD2 of BRD2.** BRD2-BD1 Y113, Y186 in BRD2-BD2 and their corresponding residue Y57 in BRD9-BD are highlighted in green. The Y113A mutation in BRD2-BD1 has been shown to disrupt acetyllysine binding.

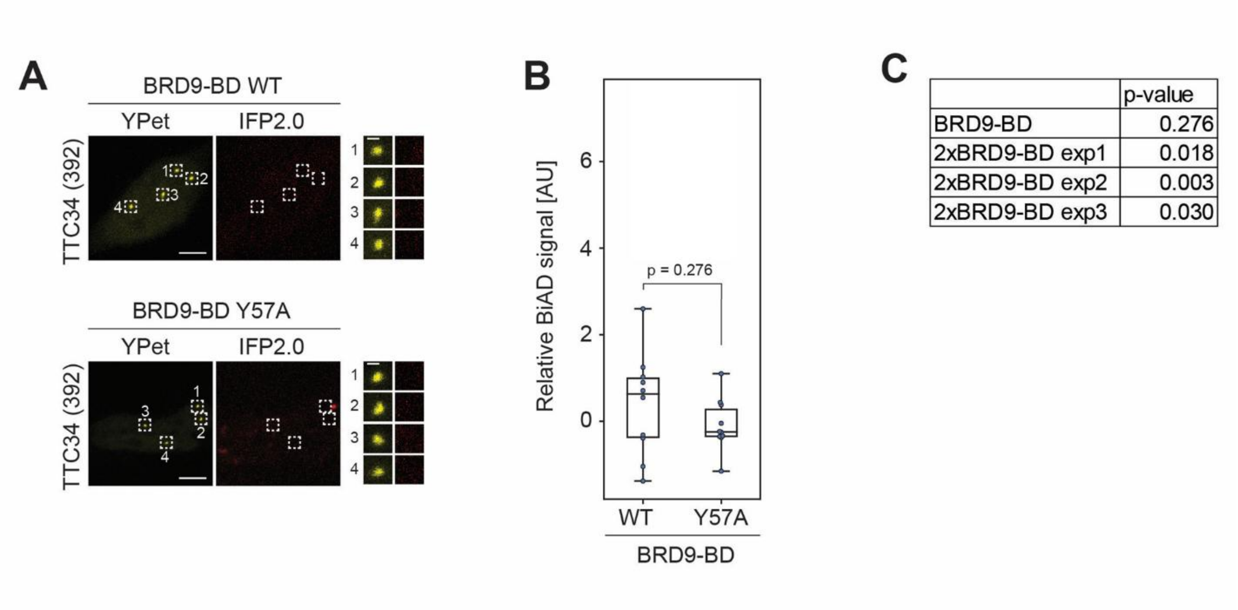


**Supplementary Figure 2: Testing of the single BRD9 bromodomain for detection of histone acetylation at the TTC34 locus.** HEK293 cells were transfected with all components of the dual-color BiAD sensor for histone acetylation detection at the TTC34 target locus with either the wildtype single BRD9BD detector (WT) or a corresponding binding-deficient mutant (Y57A). **A)** Exemplary fluorescence microscopy images showing lack of colocalization of the marker fluorophore (YPet) with a BiAD signal (IFP2.0) for both WT and Y57A detectors. Scale bars are 5 µm and 1 µm for the magnified images. **B)** Boxplot showing the relative BiAD signals of one experiment. Significance was determined via a two-tailed, unpaired t-test. **C)** P-values for differences in BiAD signal between WT and Y57A for individual experiments using either the single BRD9-BD compared with the corresponding p-values of the double domain detector 2xBRD9-BD taken form Figure 2.

| p-value | | WT | | | Y57A | | |
|---|---|---|---|---|---|---|---|
| | | DMSO | TSA 3 h | TSA 6 h | DMSO | TSA 3 h | TSA 6 h |
| WT | DMSO | 1 | 0.836 | 0.000 | 0.006 | 0.001 | 0.008 |
| | TSA 3 h | 0.836 | 1 | 0.000 | 0.018 | 0.003 | 0.023 |
| | TSA 6 h | 0.000 | 0.000 | 1 | 0.000 | 0.000 | 0.000 |
| Y57A | DMSO | 0.006 | 0.018 | 0.000 | 1 | 0.479 | 0.964 |
| | TSA 3 h | 0.001 | 0.003 | 0.000 | 0.479 | 1 | 0.461 |
| | TSA 6 h | 0.008 | 0.023 | 0.000 | 0.964 | 0.461 | 1 |

**Supplementary Figure 3: Additional information related to Figure 3.** Compilation of all pairwise p-values of the comparisons of the 2xBRD9-BD BiAD signals shown in Figure 3D. The significance of pairwise differences of BiAD signals was determined via a two-tailed, unpaired t-test. Note the highly significant signal change with BiAD sensor containing the WT 2xBRD9-BD when comparing the TSA 6 h sample, as well as the difference between WT and mutant sensors, which do not detect a signal change upon TSA treatment.
